# Development and Analytical Evaluation of a Point-of-Care Electrochemical Biosensor for Rapid and Accurate SARS-CoV-2 Detection

**DOI:** 10.1101/2023.08.08.552470

**Authors:** Mesfin Meshesha, Anik Sardar, Ruchi Supekar, Lopamudra Bhattacharjee, Soumyo Chatterjee, Nyancy Halder, Kallol Mohanta, Biplab Pal

## Abstract

The COVID-19 pandemic has underscored the critical need for rapid and accurate screening and diagnostic methods for potential respiratory viruses. Existing COVID-19 diagnostic approaches face limitations either in terms of turnaround time or accuracy. In this study, we present an electrochemical biosensor that offers nearly instantaneous and precise SARS-CoV-2 detection, suitable for point-of-care and environmental monitoring applications. The biosensor employs a stapled hACE-2 N-terminal alpha helix peptide to functionalize an in-situ grown polypyrrole conductive polymer on a nitrocellulose membrane backbone through a chemical process. We assessed the biosensor’s analytical performance using heat-inactivated omicron and delta variants of the SARS-CoV-2 virus in artificial saliva (AS) and nasal swabs (NS) samples diluted in a strong ionic solution. Virus identification was achieved through electrochemical impedance spectroscopy (EIS) and frequency analyses. The assay demonstrated a limit of detection of 40 TCID_50_/mL, with 95% sensitivity and 100% specificity. Notably, the biosensor exhibited no cross-reactivity when tested against the influenza virus. The entire testing process using the biosensor takes less than a minute. In summary, our biosensor exhibits promising potential in the battle against pandemic respiratory viruses, offering a platform for the creation of rapid, compact, portable, and point-of-care devices capable of multiplexing various viruses. This groundbreaking development has the capacity to significantly bolster our readiness and response to future viral outbreaks.

## Introduction

Recent years have witnessed an escalating global concern over respiratory virus outbreaks [1,2]. The emergence of severe acute respiratory syndrome coronavirus 2 (SARS-CoV-2) and the ensuing COVID-19 pandemic have underscored the critical need for rapid and accurate screening, diagnostics, and treatment strategies for virus-related diseases [3,4]. While significant progress has been made in developing effective COVID-19 vaccines [5], rapid and dependable virus detection remains pivotal in addressing future respiratory viral epidemic surges [6].

Various strategies have been employed for the detection of respiratory viruses, including the prominent SARS-CoV-2. These strategies majorly include nucleic acid detection methods such as reverse transcription polymerase chain reaction (RT-PCR)-based techniques and Point-of-care (PoC) lateral flow assays [7–9]. While real-time PCR stands out for its remarkable accuracy and specificity, its intricate procedure leads to delayed reporting and necessitates complex and specialized equipment and expertise [10,11]. Conversely, PoC methods such as rapid antigenic tests (RATs) offer relatively rapid outcomes, albeit at the cost of compromised accuracy [12]. In the quest for innovative approaches, the analysis of COVID-19 through exhaled breath has shown promise for self-sampling and self-testing. However, this method faces challenges such as low reproducibility and a high limit of detection (LoD), impeding its commercial viability [13].

An effective approach to mitigate such limitations can be envisaged through the development of an innovative and efficient biosensor with reliable selectivity and specific tailoring towards COVID-19 virus; in this direction, Conducting Polymer (CP)-based electrochemical biosensor is one of the promising technologies [14,15]. Conducting polymers are organic materials that are intrinsically conductive without the use of conductive fillers and owing to their elasticity, compatibility and low-cost processability, they are more preferred choice in designing and realization of biosensors [16–19]. However, construction of such biosensors based on only conducting polymers, such as, poly(acetylene), poly(3,4-ethylenedioxythiophene) (PEDOT), poly(thiophene), poly(p-phenylenevinylene) (PPV), poly(pyrrole) (PPy), poly(aniline) (PANI) and the likes, contains multiple challenges related to their amorphous nature and poor environmental stability [20,21]. Hence, development of CP based electrochemical biosensors mostly consider a composite structure comprised of porous nanostructures [22], hydrogel [23], metal nanoparticles [14], functionalized with organic and biomolecules [16], carbon composites [24], nanowire [25,26]. Generally, CPs can be mainly prepared via two approaches, namely, chemical and electrochemical methods [27]. To develop CP-based electrochemical biosensors for detecting SARS-CoV-2, immobilization of biomolecule probes, including aptamers, antibodies, ssDNA, and antigens, onto the electrode surface plays an important role in the performance of sensitivity and operational life of biosensors. Generally, these recognition elements must be directly attached to the biosensor’s surface. Physical adsorption, covalent attachment, and entrapment are the three common techniques that can be used for the immobilization of SARS-CoV-2 probes [19,28]. Among them covalent immobilization is a widely used technique where biomolecule probes are modified and functionalized by adding –NH_2_, –COOH, and other groups, and these probes are then covalently attached to functionalized monomers or CPs [16]. Shirale et. al, reported chemiresistive immunosensors based on single polypyrrole (PPy) nanowire for highly sensitive, specific, label free, and direct detection of viruses [29].

Characterization of the CP based electrochemical biosensors is mostly carried out using electrochemical impedance spectroscopy. In impedimetric biosensing applications, EIS technique is used for the detection of analytes at the sensor bio interface, and the method possess several other advantages such as simple operation, high sensitivity and short analysis time, making the process ideal for point-of-care applications [30,31]. In this direction, EIS has been employed in several biorecognition events such as lipid bilayer monitoring [32,33], DNA testing [27,34], small molecules of biological relevance [35] and cancer diagnosis [36]. In EIS, the target is recognized using antigen-antibody interaction, DNA, RNA detection, peptide-nucleic acid hybridization and aptamer-based binding. All the procedures have high sensitivity and selectivity for the target analytes and the interactions between receptor and analytes are determined by measuring potential, ion concentrations, conductance, capacitance or impedance changes [37].

This study introduces a conducting polymer-based biosensor meticulously designed for the specific detection of the SARS-CoV-2 virus. The fundamental conducting material, polypyrrole (PPy), is integrated onto a nitrocellulose membrane backbone, drawing on its established role in biosensing [38,39]. Leveraging bio-receptor interaction, we achieved selective and stable biosensor functionality by coupling lactam stapled hACE2 receptor-based peptides using a glutaraldehyde linker. A schematic illustration of the design and workflow of the biosensor development is shown in Figure 1. To assess analytical performance, the biosensor was tested using heat-inactivated omicron and delta variants of the SARS-CoV-2 virus spiked into artificial saliva and nasal swabs, the later suspended in high ionic solution. Sensitivity, specificity, and the limit of detection (LoD) were evaluated, comparing the biosensor’s performance with published rapid antigen test data. Notably, this biosensor incorporates a robust electrical signal data analysis strategy, meticulously optimizing the frequency range in impedance measurement. This strategic refinement effectively neutralizes the influence of noise signals that could compromise result accuracy. This algorithmic augmentation ensures precise and reliable virus detection, thereby enhancing the overall efficacy and durability of the biosensor system.

**Figure 1.**
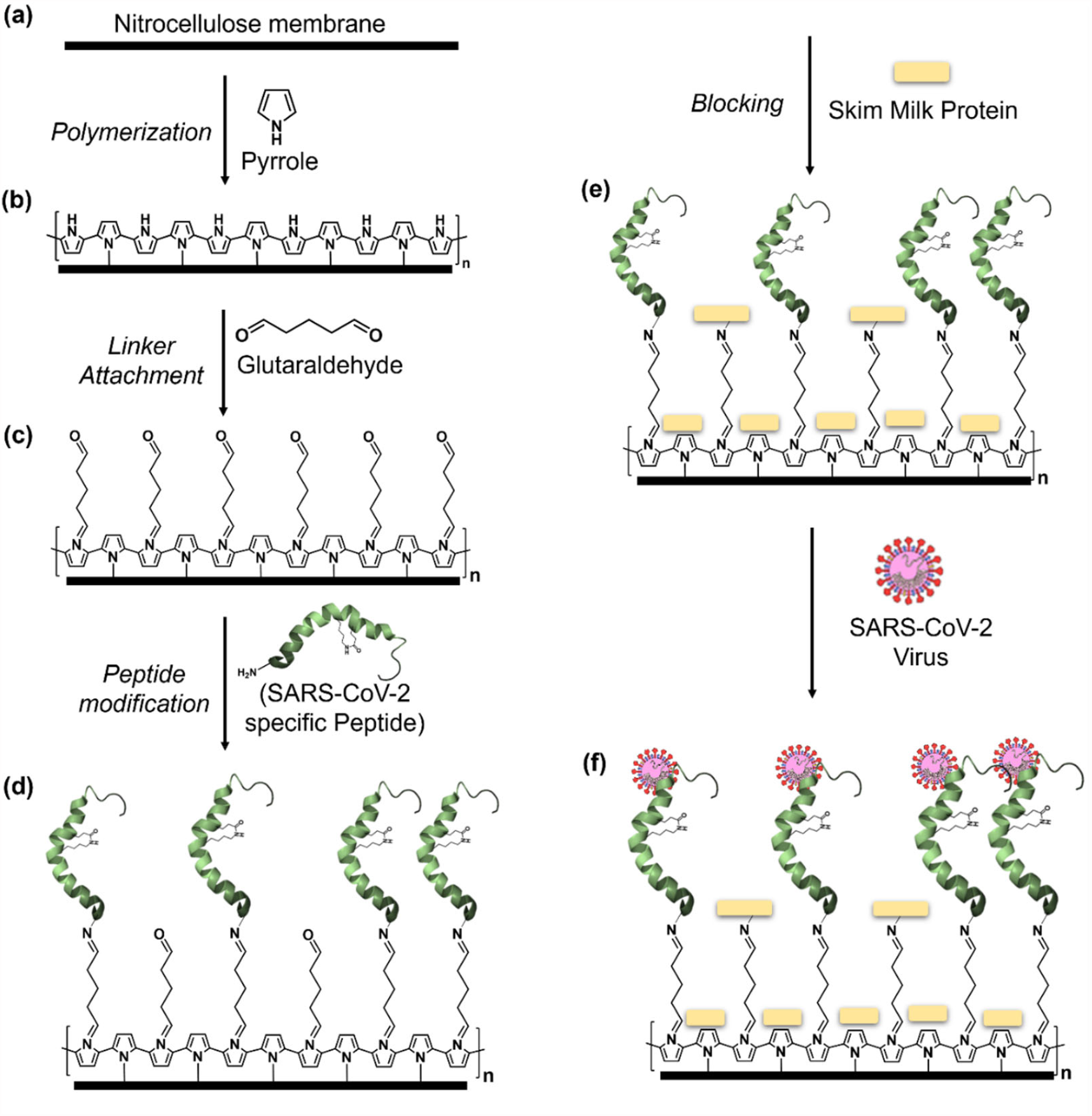
Schematic presentation of biosensor development. (a) Nitrocellulose membrane (NC) (1mm × 10mm in dimension) as the base of the sensor substrate, (b) polymerization of conducting polymer (polypyrrole) on NC membrane, (c) covalent attachment of organic linker (glutaraldehyde), (d) functionalization with lactam stapled SARS-CoV-2 specific peptide, (e) blocking with skim milk protein and (f) interaction of the SARS-CoV-2 virus with the receptor peptide.

## Materials and Methods

### Reagents

Pyrrole (C_4_H_5_N) 98.0% (Merck, Germany) was vacuum distilled before use. Ammonium persulphate ((NH_4_)_2_S_2_O_8_) 98.0% (APS) and glutaraldehyde (OHC(CH_2_)_3_CHO) (Grade 1, 25% in water) were obtained from Merck, Germany. Nitrocellulose membrane (pore size 0.45 μm) was purchased from Fisher Scientific. All other chemicals were of analytical grade and used without further purification. Hydrochloric acid (HCl), phosphate buffer saline (PBS. pH 7.4), and de-ionized water were obtained from Merck (Germany), SRL (India), and Emplura (Merck, Germany), respectively.

The hACE2 peptide sequence (Figure S1) used in this study was adopted from Maas et.al [40] and was purchased from GL Biochem (Shanghai) Ltd. The peptide sequence specifically binds to the receptor binding domain (RBD) region of the SARS-CoV-2 spike protein. The lactam-linked stapling modification on the peptide at position K12 and E20 (i, i+4) enhances the stability of its three-dimensional structure. For the peptide-virus binding evaluation experiments, the peptide was tagged with the Alexa Fluor 488 NHS ester (green) dye (A270022, antibodies.com, UK).

### Viruses

Viruses utilized in this study were SARS-CoV-2 lineage B.1.617.2 (Delta Variant) culture fluid (heat inactivated, 0810624CFHI, Zeptometrix LLC, USA) and SARS-CoV-2 lineage B.1.1.529 (Omicron Variant) culture fluid (UV inactivated, 0810642UV, Zeptometrix LLC, USA) and (heat inactivated, 0810642CFHI, Zeptometrix LLC, USA). For control experiments, Influenza vaccine (Fluarix-Tetra 2021 South, GlaxoSmithKline Biologicals, Germany) containing attenuated mix of Influenza A and B viruses (A/Victoria/2570/2019 (H1N1), A/Hong Kong/2671/2019 (H3N2), B/Washington/02/2019 and B/Phuket/3037/2013) was used. We directly employed the pre-inactivation viral titers provided by the manufacturer in TCID_50_/mL, which can be found in the product inserts for detailed reference. Additionally, we quantified the virus stocks through RT-PCR analysis using standard curve generated with the 10-fold serially diluted positive control (provided with the kit) of known copy number using the Coronavirus COVID-19 Genesig real-time PCR assay kit (Z-Path-COVID-19-CE, Primer Design Ltd., UK) (Figure S2). To evaluate the sensitivity and specificity of the biosensor, the virus, originally present in culture supernatant, was subjected to buffer exchange into the desired media (Artificial saliva, SAE0149, Sigma-Aldrich, USA). The buffer exchange process was conducted using Nanosep 3K Omega columns (OD003C33, Pall corporations). During the experimental procedure of detecting viruses, 1 μL of freshly prepared viral solution was dropped on the biosensor using micropipette and impedance measurement was conducted.

### Development of the biosensors

Development of the biosensors on the nitrocellulose substrate was carried out following a sequential approach as shown in Figure 1. Initially, specific dimensions of nitrocellulose membranes (1 mm × 10 mm) were coated with polypyrrole via oxidative polymerization of pyrrole monomer in an acidic medium using ammonium persulfate (APS) as the oxidizing agent. A mixture of 0.1 M pyrrole and 0.1 M APS in 1 M HCl was used, and the substrates were immersed in this solution (1:1 ratio) at 10°C for *in situ* polypyrrole deposition, which took around 120 minutes. The resulting polypyrrole-coated substrates were rinsed with deionized water and air-dried. Following substrate preparation, a linker, glutaraldehyde, was covalently attached by treating the polypyrrole-coated substrates with 25% glutaraldehyde for 4 hours, facilitating flexible bridges for effective binding of biomolecules [41,42]. After washing with phosphate-buffered saline (PBS), site-specific SARS-CoV-2 RBD lactam-stapled hACE-2 peptide was covalently immobilized onto the glutaraldehyde-treated substrates at 30 μg/ml concentration in 1X PBS for 16 hours. Excess and unbound peptide were removed by washing with 1X Tris-buffered saline with Tween20 (TBST). To reduce non-specific binding, 5% skim milk in 1X PBS was used to block binding sites for 90 minutes. After further washing with 1X TBST and deionized water, the substrates were vacuum-dried for about 2 hours, serving as the basis for biosensor fabrication.

### Impedance measurement

In the experimental phase, the sensor strips are affixed to electrical connectors at both ends using silver paste. These connected sensors are then inserted into an impedance analyzer, specifically the PGSTAT204 potentiostat from Autolab (Germany), for virus detection experiments. As a two-electrode sensor configuration is employed, the potentiostat’s electrode arrangement adheres to the designated scheme (Figure S3), involving a working electrode biased relative to the reference electrode without a ground connection, while the source and counter electrodes serve to facilitate charge flow. An alternating current (AC) bias of 100 mV is applied, and the frequency is scanned logarithmically from 20 kHz to 40 kHz with 10 points per decade. The initial step entails measuring the impedance of a blank sensor. Subsequently, 1 μL of either virus or control solution sample is dispensed onto the sensor’s central region. Within 5-10 seconds of sample exposure, another impedance measurement is initiated. The first scan of the sensor (either blank or dry) is denoted as the initial impedance, while the scan following sample exposure is termed the final impedance. The entire frequency scan duration for each set of measurements (initial and final impedance) is approximately 20 seconds. Given that significant impedance changes are primarily observed in the magnitude upon sample exposure, the analysis focuses solely on the magnitude of impedance. To account for potential variations in the initial impedance values among sensors from different batches, a normalized impedance change parameter (dZ/Z) is introduced using the formula:

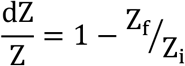

where dZ represents the change in total impedance post-sample exposure, Z_f_ denotes the final impedance after one minute of sample exposure, and Z_i_ signifies the initial impedance of the sensor. Notably, this parameter’s value is also influenced by the probing frequency. The impedance analysis is carried out within a specific frequency range of 1 kHz to 100 kHz, chosen and fine-tuned through empirical optimization.

## Results and Discussion

The biosensor was functionalized with a specific lactam-linked stapled hACE-2 N-terminal alpha helix peptide for SARS-CoV-2 recognition. The peptide was chosen based on literature describing its interaction with the SARS-CoV-2 spike protein’s receptor binding site (RBD) and hACE-2 transmembrane protease domain, facilitating viral fusion and cellular entry [43–45]. Engineered hACE-2 transmembrane domains were investigated to obstruct the RBD-hACE-2 interaction for potential therapeutics. However, alpha 1-helix-based hACE2 peptides lose bioactivity in solution, affecting RBD binding [46,47]. To address this issue, a lactam i, i+4 stapling modification was utilized to stabilize the structure of the hACE-2 peptide [40,48].

In our investigation, we employed the 35-mer lactam-based stapled hACE2 N-terminal α1-helix inhibitor 1 sequence as outlined by Maas et al to functionalize our conductive polymer biosensor, with the aim of establishing a rapid SARS-CoV-2 detection method [40]. To evaluate the virus’s selective binding to this peptide, we initiated the process by coating a glass slide with PPy, introducing a glutaraldehyde linker, and functionalizing the biosensor with the stapled hACE-2 peptide at a concentration of 30 μg/mL. Subsequently, a 5% skim milk protein solution was used to block non-specific binding on the sensor. Next, the SARS-CoV-2 Omicron variant virus, at a concentration of 3 log_10_TCID_50_/mL in artificial saliva, was applied to the biosensor and incubated for one hour at room temperature. Following a washing step, the biosensor was exposed to an Alexa 488-tagged hACE-2 peptide to interact with the virus-biosensor complex. After additional washing with 1X TBST, images were captured via fluorescence microscopy. To ascertain specificity, a heat-inactivated influenza virus was used as a control. As anticipated, the biosensor demonstrated selective binding to the SARS-CoV-2 virus, exhibiting no binding to the Influenza viruses, even at a high concentration of 120 μg/mL of haemagglutinin (HA) protein thereby affirming no cross reactivity (Figure 2).

**Figure 2.**
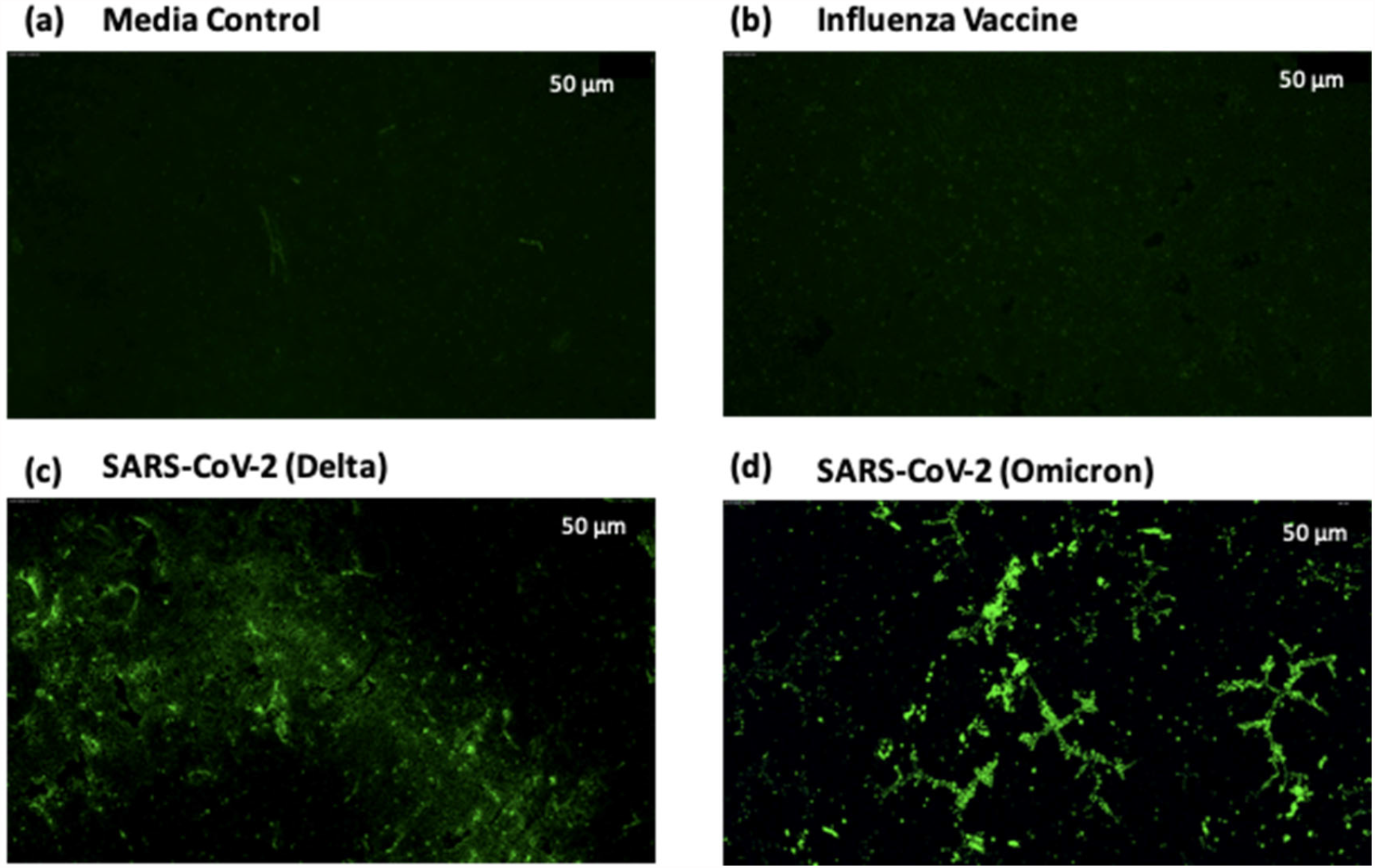
Selective binding of SARS-CoV-2 to a lactam stapled hACE-2 functionalized on PPy substrate on glass slide. The PPy coated and glutaraldehyde linked glass slides were treated with hACE-2 peptide, blocked with skim milk protein before addition of virus or controls. Alexa Fluor 488 fluorophore tagged peptide was then used to probe virus binding. (a) Artificial saliva without virus spike-in was used as a media control. (b) Heat attenuated influenza vaccine containing a mix of Influenza A (H1N1, H3N2) and B viruses. (c) SARS-CoV-2 delta variant at concentration of 10^5^ virus copies/μL (d) SARS-CoV-2 omicron variant at concentration of 10^5^ virus copies/μL

### Characterization of biosensor and detection of SARS-CoV-2 using electrochemical impedance spectroscopy

As previously mentioned, EIS is a widely recognized technique for characterizing impedimetric biosensors and analyzing interfacial properties associated with bio-recognition events [49]. In line with EIS’s suitability, we conducted impedance measurements (details provided in the materials and methods section) at various stages of sensor development to ensure fabrication process reproducibility and establish baseline impedance levels for each biosensor development step. As depicted in Figure 3a, distinct bands of absolute impedance were evident after addition of each component on the nitrocellulose membrane substrate, with steadily increasing impedance. Polypyrrole functions as a *p*-type semiconductor where holes are the dominant charge carriers [50]. The incremental impedance rise suggests a gradual depletion of charge carriers due to the negative charges of entities such as glutaraldehyde, peptides, and skim milk protein binding to PPy. These negative entities attract positive holes from the conducting polymer backbone, diminishing the carrier count and thus elevating total impedance [51].

**Figure 3.**
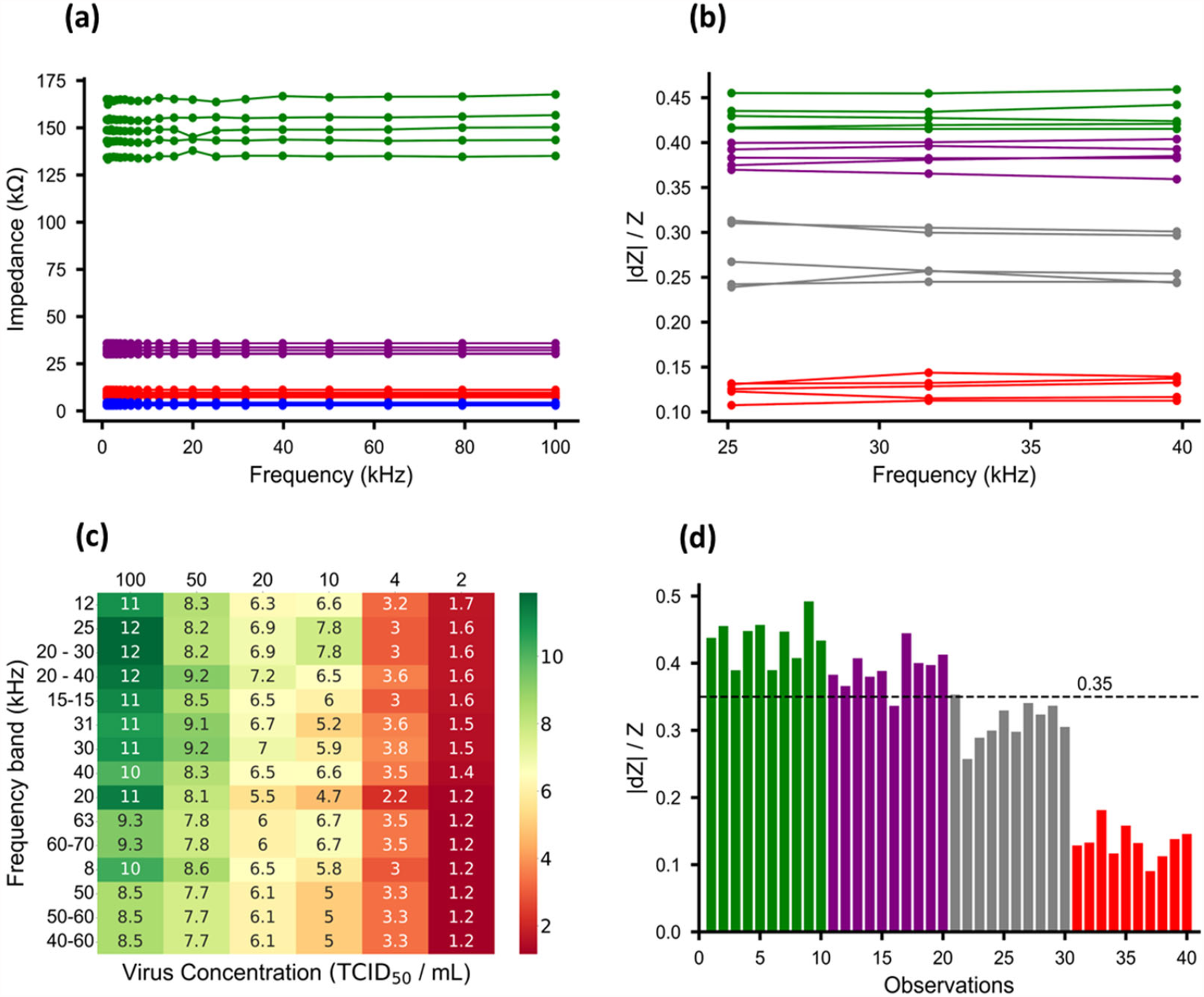
Characterization of biosensor and detection of SARS-CoV-2 virus using electrochemical impedance spectroscopy. (a) Characteristic impedance measurement at different stages of sensor development across a range of frequencies. Blue lines: after coating with polypyrrole, Red lines: after GA linker addition, Purple lines: after hACE2 peptide attachment, and Green lines: after blocking with skimmed milk protein. (b) Sensor response in terms of relative impedance change (|dZ|/Z) for different virus concentrations and media control. Green lines: media control, Purple, Grey and Red lines are virus in artificial saliva at concentrations of 20, 40 and 1000 TCID_50_/mL, respectively. (c) heatmap for optimization of separation factor between virus and control class. Frequency band corresponds to the optimum separation factor (Sf) between media control and different virus concentrations. The greener shades indicate higher Sf values while the reddish shades indicate lower Sf values. The rows are sorted in descending order based on the Sf values of viral RNA copies 20 TCID_50_/mL, and (d) Sensor response in terms of relative impedance change (|dZ|/Z) for different virus concentration and media control (10 representative samples from 20 replicates) separated by a threshold line. Green bars: media control, purple, grey and red bars are viral RNA copies (20 TCID_50_/mL, 40 TCID_50_/mL and 1000 TCID_50_/mL respectively). A dashed black threshold line is drawn at 3-standard deviations below the mean of the control data set.

Subsequently, impedance alterations were gauged upon the introduction of the quantified virus. Specifically, 1 μL of SARS-CoV-2 virus was added to the biosensor in five independent replicates, with high concentration (1000 TCID_50_/mL) and two lower concentrations (40 and 20 TCID_50_/mL), and impedance measurements were conducted (Figure 3b). As a negative control, 1 μL of artificial saliva was applied to offset impedance shifts arising from charges within the virus media. The outcomes unveiled distinctive impedance bands corresponding to virus concentration on the biosensor, with the most pronounced distinction between virus data (highlighted by red lines in Figure 3b) and control data (emphasized by green lines in Figure 3b) evident at the highest virus concentration (1000 TCID_50_/mL). Nevertheless, the data also exhibited fluctuations and noise, particularly within the lower frequency ranges.

In order to mitigate the impact of noise, we embarked on data pre-processing strategies with the goal of identifying the optimal frequency or frequency range that maximizes the discernment between the control and virus classes, particularly for lower virus concentrations such as 20 and 40 TCID_50_/mL. Initially, the sensor’s responses to virus and control samples were recorded in terms of the modulus of impedance change, represented by the alteration in impedance (dZ) from its initial value (Z) before the introduction of the virus or control droplet. Subsequently, segregation of control data from various virus concentration data was achieved using the 3-sigma rule principle [52], assuming normal data distribution. Figures S4a and b in the supplementary information showcase the characteristic normal distribution for the 40 TCID_50_/mL virus concentration, alongside the results of normality tests. Leveraging the assumed normal distribution of other data points, we computed the separation factor (Sf) between the virus and control classes using the following formula,

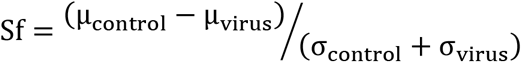

where, Sf denotes separation factor, μ_control_ and μ_virus_ represent the means of the control and virus data, respectively, while σ_control_ and σ_virus_ represent their respective standard deviations.

The separation factor between the control and varying virus concentrations was computed across individual frequencies and a range of frequency bands. Subsequently, frequencies were sorted based on descending sf values specifically for the lowest virus concentration (20 TCID_50_/mL), leading to the creation of a heatmap (Figure 3c). A higher separation factor denotes a more pronounced distinction between the virus and control classes. In this study, a separation factor of 3 indicates strong capability, signifying a high likelihood of distinguishing the virus from the control or an error probability of 0.27% in virus detection. Similarly, a separation factor of 1.96 serves as a cutoff, representing a 95% probability of separation or a 5% error probability in virus detection. As shown in Figure 3c, Sf values are depicted for both control and diverse virus concentrations. Notably, the heatmap highlights a peak separation factor at 12kHz for the 20 TCID_50_/mL virus concentration. However, within the 20kHz to 40kHz frequency band, consistently higher separation factors are evident across most virus concentrations. This initial analysis was used to determine the preliminary limit of detection of the biosensor (see detail below).

### Determination of virus detection based on relative impedance threshold

In the quest to determine the classification of individual sample tests as positive or negative virus detection, we devised a comprehensive threshold line computation strategy encompassing distinct steps. First, the optimal input frequency was pinpointed as outlined previously. Subsequently, the central tendency of the relative impedance (dZ/Z) values from control data corresponding to the chosen frequency band for each test data was calculated, utilizing the median as the central tendency metric. The mean (μ_control_) and standard deviation (σ_control_) were then computed across all control test data. Then, a decisive threshold line was established based on the following the equation:

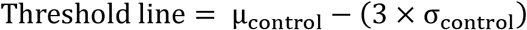

Responses below this established threshold line were identified as belonging to the virus class, whereas those exceeding it were categorized as control class. The specified threshold, set at 0.35, notably exhibited exceptional sensitivity of 95% in distinguishing the 4 TCID_50_/mL virus concentration data from control data, as shown in Figure 3d (with 10 representative samples from each virus concentration among the 20 independent replicates). Alternatively, a machine learning (ML) approach was adopted to determine the threshold value. Following a parallel methodology, an optimal frequency range was selected for data processing, with |dZ|/Z values serving as pivotal feature variables for the model. The dataset, constituting 40 test data from both control (20) and virus experiments (20), was randomly divided into a 75:25 ratio to form distinct training and testing datasets. A decision tree model of depth 1 was fitted, culminating in a threshold value of approximately 0.37 (Figure S5). This threshold was subsequently tested on the designated dataset, yielding an impressive 100% accuracy in discerning 40 TCID_50_/mL virus concentration data from control data. Furthermore, the established threshold from the decision tree model was rigorously validated using a completely new dataset comprising 25 virus test samples and 25 control test samples, yielding remarkable outcomes of 100% sensitivity. Detailed validation results are presented in the supplementary figure (Figure S5). However, in adherence to a conservative approach and due to the limited dataset for the machine learning model, we retained the earlier formula for drawing threshold lines.

### Limit of detection, Sensitivity, and specificity of the biosensor

To comprehensively evaluate our biosensor’s analytical performance, we conducted tests to establish the limit of detection (LoD), sensitivity, and specificity. Initially, we performed a 10-fold assessment to determine preliminary LoD (Figure S6) followed by testing selected concentrations from a 2-fold serially diluted virus analyte in artificial saliva (Figure 4a). Integrating these tests with our previous analyses (refer to Figures 3b and 3d), we identified the concentration yielding a minimal yet above the cut-off separation factor (1.96) as the preliminary LoD. Consequently, we set the assay’s detection limit at 40 TCID_50_/mL virus concentration, affirming this determination through 20 replicates at the specified LoD, in accordance with the requirements of Emergency Use Authorization program of Food and Drug Administration (FDA EUA), aiming for 95% sensitivity. In parallel, 20 replicates of artificial saliva without virus were analyzed. As depicted in Figure 4b, the assay exhibited remarkable sensitivity at this LoD, detecting 19 positives out of 20 replicates and accurately identifying all negatives, providing 95% sensitivity and 100% specificity. Furthermore, considering that many COVID tests utilize nasal swab specimens, we acquired nasal swabs from healthy donors suspended in a high ionic buffer (0.45 M KCl) and spiked with equal virus concentration (40 TCID_50_/mL) for assessment with our biosensor. Notably, the results showcased similar sensitivity to that observed with virus in artificial saliva (Figure 4c). There was no notable difference in sensitivity between omicron and delta variants of SARS-CoV-2 in sensitivity (Figure S8). Furthermore, the biosensor was tested with heat attenuated influenza vaccine containing a mix of Influenza A (H1N1, H3N2) and B viruses and did not show any cross reactivity (Figure 4d).

**Figure 4.**
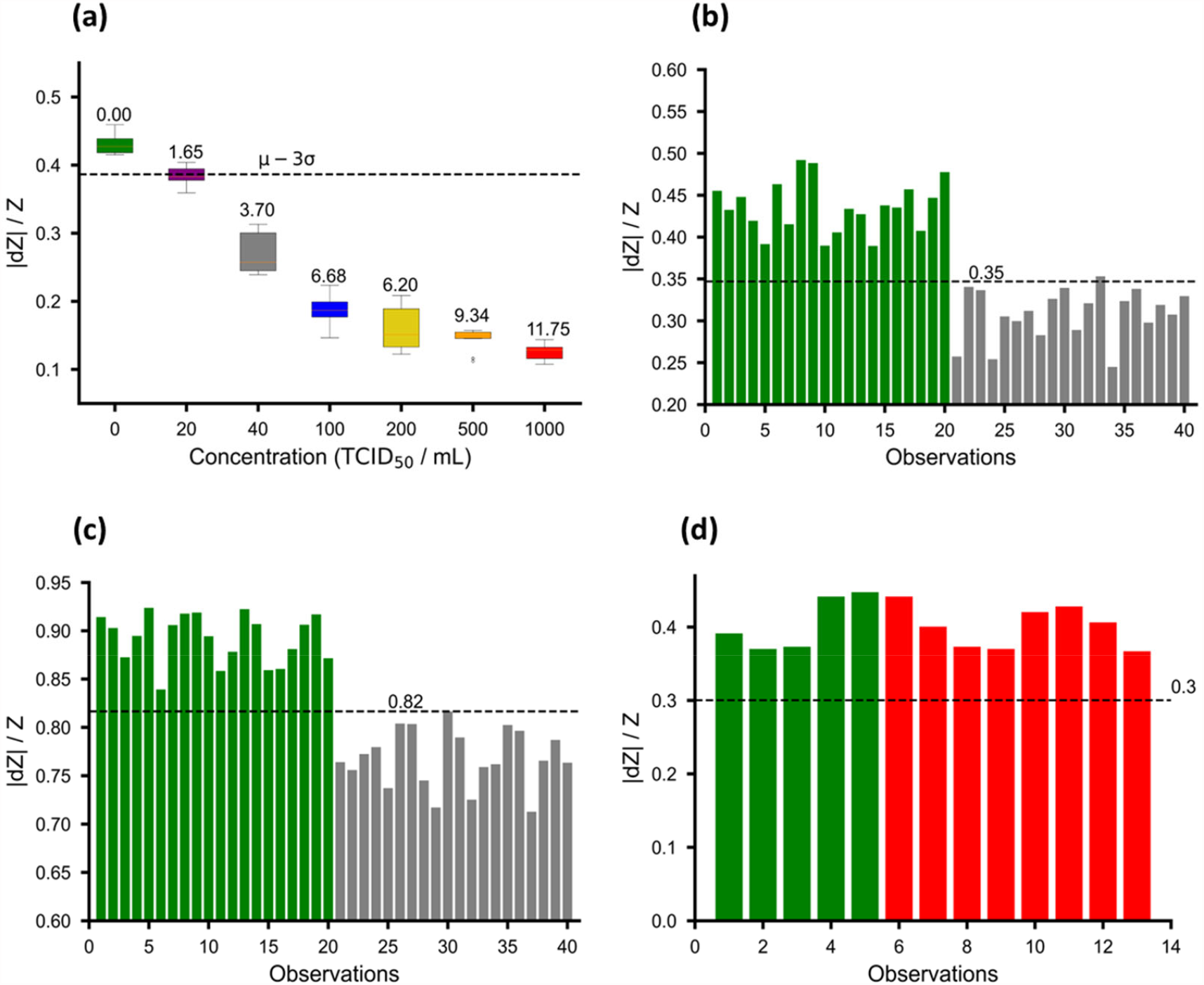
Classification of virus detection: (a) Relative impedance change (|dZ|/Z) for different virus concentrations and media control. The colors indicate virus concentrations: green (media control), purple (20 TCID_50_/mL), grey (40 TCID_50_/mL), blue (10 TCID_50_/mL), yellow (20 TCID_50_/mL), orange (50 TCID_50_/mL), and red (1000 TCID_50_/mL). Sf values are annotated for each box. (b) Limit of detection validation at 40 TCID_50_/mL. Y-axis: relative impedance value (|dZ|/Z). Green and grey bars represent media control and virus, respectively. (c) Comparable sensitivity of virus spiked in nasal swabs in 0.45M KCl buffer. Y-axis: relative impedance value (|dZ|/Z). Green and grey bars represent media control and virus, respectively. (d) Relative impedance changes for influenza vaccine and media control. y-axis: relative impedance value (|dZ|/Z). Green and grey bars represent media control and influenza, respectively. Threshold lines are denoted by black dashed lines for all figures and represent 3-standard deviation below the mean of control data’s relative impedance change.

The determined 95% LoD threshold of our assay is 40 TCID_50_/mL as determined by the manufacturer’s pre-inactivation analysis. To facilitate fair comparisons with other studies evaluating RAT analytical performance, we determined viral RNA copies corresponding to dilutions of TCID_50_/mL concentrations provided by the manufacturer using RT-PCR (Figure S7). Based on this analysis our 95% LoD threshold corresponds to 6.6 log10 RNA copies/mL. Studies on the analytical performance of RATs are diverse due to differences in study design and methods, sample size, and type of RAT evaluated which demonstrated variable performance, and sometimes with reports of contradicting results for similar assays making comparisons difficult [53–55]. Furthermore, variability in viral quantification, as TCID_50_, plaque forming units (pfu) and RT-PCR based viral RNA copies introduce bias to make comparisons [56]. However, within this context and limitations, our biosensor demonstrates comparable or better analytical performance with commonly used rapid antigen tests. For example, Corman et al. evaluated 95% LoD of seven antigen point-of-care tests using 138 clinical specimens and reported LoDs ranging between 6.32 log_10_ and 7.46 log_10_ RNA copies per swab [57]. Similarly, Deerain et al. [55] evaluated the analytical sensitivity of commercially available 10 RATs using representative Delta and Omicron isolates cultured from clinical samples and reported 95% LoDs of 6.50 log_10_ copies/mL and the Omicron variant at 6.39 log_10_ copies/mL for Delta and Omicron variants respectively. Put together, our data indicate that our Biosensor has comparable, in most cases better analytical performance with commercially available RATs. While our developed Biosensor like commercially available rapid antigen tests may not match the sensitivity of reverse transcription-PCR (RT-PCR) tests, their distinct advantage lies in their rapid detection capabilities for individuals with high viral loads that drive transmission. This attribute makes our biosensor a valuable tool for clinical and public health applications, particularly in various settings where swift and efficient detection is essential.

Our biosensor offers a distinct advantage over Rapid Antigen Tests due to heightened sensitivity achieved through miniaturization. Currently at 1 mm x 10 mm, it can be further reduced to 50 μm x 25 μm via photolithography, enhancing sensitivity by minimizing the conducting surface area, thus detecting extremely low virus concentrations, and improving its Limit of Detection (LoD). This sensitivity improvement is rooted in the reduced dimensions of the active binding region [58], optimizing portability, sensitivity, and overall performance while reducing power consumption compared to conventional designs [59]. Miniaturization also facilitates more efficient mass transport, faster analyte binding, superior multiplexing capabilities for various applications [60], yet challenges like functionalization and parallel reading of closely spaced sensor locations must be tackled [61].

## Conclusions

In summary, our study has successfully developed a rapid biosensor tailored for precise SARS-CoV-2 detection. This was achieved by functionalizing a conductive polymer biosensor with a lactam stapled hACE-2 N-terminal alpha helix peptide, endowing it with exceptional selectivity. This selectivity was corroborated through both fluorescence microscopy and impedance measurement, enabling accurate virus detection in minuscule volumes of artificial saliva and nasal swabs. The biosensor demonstrated robust analytical performance, boasting 95% sensitivity and 100% specificity at a LoD of 40 TCID_50_/mL. These results are comparable to, and in many cases superior to, commercial rapid antigen tests. Subsequent clinical evaluation is necessary to assess its practical utility. Furthermore, our biosensor showcased rapid and precise SARS-CoV-2 detection, yielding results in under a minute. A noteworthy advantage of our biosensor lies in its potential for heightened sensitivity through miniaturization. This enhancement translates into improved portability for point-of-care applications, reduced power consumption, accelerated response times, and the capability to simultaneously detect multiple analytes. These attributes render our biosensor suitable for diverse applications in medical diagnostics and environmental monitoring.

## Author Contributions

Conceptualization: Biplab Pal, Mesfin Meshesha

Data curation: Soumyo Chatterjee, Anik Sardar, Kallol Mohanta.

Formal analysis and algorithm development: Anik Sardar, Biplab Pal, Mesfin Meshesha, Kallol Mohanta

Investigation: Kallol Mohanta, Soumyo Chatterjee, Lopamudra Bhattacharjee, Nyancy Halder.

Methodology: Mesfin Meshesha, Lopamudra Bhattacharjee, Nyancy Halder, Kallol Mohanta, Soumyo Chatterjee, Ruchi Supekar,

Project administration: Biplab Pal, Mesfin Meshesha

Supervision: Mesfin Meshesha, Biplab Pal

Writing – original draft: Mesfin Meshesha, Anik Sardar, Lopamudra Bhattacharjee, Nyancy Halder, Soumyo Chatterjee, Kallol Mohanta, Ruchi Supekar,

Writing – review & editing: Mesfin Meshesha

All authors have read and approved the manuscript.

## Acknowledgement

The authors thank Mr. Conrad Bessemer, Chairman and co-founder, Opteev Technologies Inc, for providing financial support for the entire work. The authors also thank Indian Institute of Technology, Kharagpur and International Institute of Innovation and Technology, Kolkata for providing research facilities. Ms. Ankita Saha, Mr. Shaunak Guha and Ms. Sreyashi Paul are acknowledged for their assistance in lab work. Mr. Biswadeep Ghoshal and Ms. Jayabani Ghosh are acknowledged for their support in sensor data analysis.

## References

1. Dhama, K.; Khan, S.; Tiwari, R.; Sircar, S.; Bhat, S.; Malik, Y.S.; Singh, K.P.; Chaicumpa, W.; Bonilla-Aldana, D.K.; Rodriguez-Morales, A.J. Coronavirus Disease 2019–COVID-19. Clin. Microbiol. Rev. 2020, 33, e00028–20, doi:10.1128/CMR.00028-20.

2. Pascarella, G.; Strumia, A.; Piliego, C.; Bruno, F.; Del Buono, R.; Costa, F.; Scarlata, S.; Agrò, F.E. COVID-19 Diagnosis and Management: A Comprehensive Review. J. Intern. Med. 2020, 288, 192–206, doi:10.1111/joim.13091.

3. More, N.; Ranglani, D.; Kharche, S.; Choppadandi, M.; Ghosh, S.; Vaidya, S.; Kapusetti, G. Current Challenges in Identification of Clinical Characteristics and Detection of COVID-19: A Comprehensive Review. Meas. Sens. 2021, 16, 100052, doi:10.1016/j.measen.2021.100052.

4. Mohite, V.; Vyas, K.; Phadke, G.; Rawtani, D. Challenges and Future Aspects of COVID-19 Monitoring and Detection. In COVID-19 in the Environment; Elsevier, 2022; pp. 131–150 ISBN 978-0-323-90272-4.

5. Rahman, Md.M.; Masum, Md.H.U.; Wajed, S.; Talukder, A. A Comprehensive Review on COVID-19 Vaccines: Development, Effectiveness, Adverse Effects, Distribution and Challenges. VirusDisease 2022, 33, 1–22, doi:10.1007/s13337-022-00755-1.

6. Ndwandwe, D.; Wiysonge, C.S. COVID-19 Vaccines. Curr. Opin. Immunol. 2021, 71, 111–116, doi:10.1016/j.coi.2021.07.003.

7. Pokhrel, P.; Hu, C.; Mao, H. Detecting the Coronavirus (COVID-19). ACS Sens. 2020, 5, 2283–2296, doi:10.1021/acssensors.0c01153.

8. Ji, T.; Liu, Z.; Wang, G.; Guo, X.; Akbar Khan, S.; Lai, C.; Chen, H.; Huang, S.; Xia, S.; Chen, B.; et al. Detection of COVID-19: A Review of the Current Literature and Future Perspectives. Biosens. Bioelectron. 2020, 166, 112455, doi:10.1016/j.bios.2020.112455.

9. Filchakova, O.; Dossym, D.; Ilyas, A.; Kuanysheva, T.; Abdizhamil, A.; Bukasov, R. Review of COVID-19 Testing and Diagnostic Methods. Talanta 2022, 244, 123409, doi:10.1016/j.talanta.2022.123409.

10. Teymouri, M.; Mollazadeh, S.; Mortazavi, H.; Naderi Ghale-noie, Z.; Keyvani, V.; Aghababaei, F.; Hamblin, M. R.; Abbaszadeh-Goudarzi, G.; Pourghadamyari, H.; Hashemian, S.M.R.; et al. Recent Advances and Challenges of RT-PCR Tests for the Diagnosis of COVID-19. Pathol. - Res. Pract. 2021, 221, 153443, doi:10.1016/j.prp.2021.153443.

11. Feng, W.; Newbigging, A.M.; Le, C.; Pang, B.; Peng, H.; Cao, Y.; Wu, J.; Abbas, G.; Song, J.; Wang, D.-B.; et al. Molecular Diagnosis of COVID-19: Challenges and Research Needs. Anal. Chem. 2020, 92, 10196–10209, doi:10.1021/acs.analchem.0c02060.

12. Routsias, J.G.; Mavrouli, M.; Tsoplou, P.; Dioikitopoulou, K.; Tsakris, A. Diagnostic Performance of Rapid Antigen Tests (RATs) for SARS-CoV-2 and Their Efficacy in Monitoring the Infectiousness of COVID-19 Patients. Sci. Rep. 2021, 11, 22863, doi:10.1038/s41598-021-02197-z.

13. Giovannini, G.; Haick, H.; Garoli, D. Detecting COVID-19 from Breath: A Game Changer for a Big Challenge. ACS Sens. 2021, 6, 1408–1417, doi:10.1021/acssensors.1c00312.

14. Kwon, O.S.; Park, S.J.; Lee, J.S.; Park, E.; Kim, T.; Park, H.-W.; You, S.A.; Yoon, H.; Jang, J. Multidimensional Conducting Polymer Nanotubes for Ultrasensitive Chemical Nerve Agent Sensing. Nano Lett. 2012, 12, 2797–2802, doi:10.1021/nl204587t.

15. Kangkamano, T.; Numnuam, A.; Limbut, W.; Kanatharana, P.; Vilaivan, T.; Thavarungkul, P. Pyrrolidinyl PNA Polypyrrole/Silver Nanofoam Electrode as a Novel Label-Free Electrochemical MiRNA-21 Biosensor. Biosens. Bioelectron. 2018, 102, 217–225, doi:10.1016/j.bios.2017.11.024.

16. Tran, V.V.; Tran, N.H.T.; Hwang, H.S.; Chang, M. Development Strategies of Conducting Polymer-Based Electrochemical Biosensors for Virus Biomarkers: Potential for Rapid COVID-19 Detection. Biosens. Bioelectron. 2021, 182, 113192, doi:10.1016/j.bios.2021.113192.

17. Gerard, M. Application of Conducting Polymers to Biosensors. Biosens. Bioelectron. 2002, 17, 345–359, doi:10.1016/S0956-5663(01)00312-8.

18. Contractor, A.Q.; Sureshkumar, T.N.; Narayanan, R.; Sukeerthi, S.; Lal, R.; Srinivasa, R.S. Conducting Polymer-Based Biosensors. Electrochimica Acta 1994, 39, 1321–1324, doi:10.1016/0013-4686(94)E0054-4.

19. Lakard, B. Electrochemical Biosensors Based on Conducting Polymers: A Review. Appl. Sci. 2020, 10, 6614, doi:10.3390/app10186614.

20. Yoon, H.; Jang, J. Conducting-Polymer Nanomaterials for High-Performance Sensor Applications: Issues and Challenges. Adv. Funct. Mater. 2009, 19, 1567–1576, doi:10.1002/adfm.200801141.

21. Omastová, M.; Kosina, S.; Pionteck, J.; Janke, A.; Pavlinec, J. Electrical Properties and Stability of Polypyrrole Containing Conducting Polymer Composites. Synth. Met. 1996, 81, 49–57, doi:10.1016/0379-6779(96)80228-1.

22. Song, Z.; Ma, Y.; Morrin, A.; Ding, C.; Luo, X. Preparation and Electrochemical Sensing Application of Porous Conducting Polymers. TrAC Trends Anal. Chem. 2021, 135, 116155, doi:10.1016/j.trac.2020.116155.

23. John, A.; Benny, L.; Cherian, A.R.; Narahari, S.Y.; Varghese, A.; Hegde, G. Electrochemical Sensors Using Conducting Polymer/Noble Metal Nanoparticle Nanocomposites for the Detection of Various Analytes: A Review. J. Nanostructure Chem. 2021, 11, 1–31, doi:10.1007/s40097-020-00372-8.

24. Fahmy, H.M.; Abu Serea, E.S.; Salah-Eldin, R.E.; Al-Hafiry, S.A.; Ali, M.K.; Shalan, A.E.; Lanceros-Méndez, S. Recent Progress in Graphene- and Related Carbon-Nanomaterial-Based Electrochemical Biosensors for Early Disease Detection. ACS Biomater. Sci. Eng. 2022, 8, 964–1000, doi:10.1021/acsbiomaterials.1c00710.

25. Bangar, M.A.; Shirale, D.J.; Chen, W.; Myung, N.V.; Mulchandani, A. Single Conducting Polymer Nanowire Chemiresistive Label-Free Immunosensor for Cancer Biomarker. Anal. Chem. 2009, 81, 2168–2175, doi:10.1021/ac802319f.

26. Chartuprayoon, N.; Rheem, Y.; Ng, J.C.K.; Nam, J.; Chen, W.; Myung, N.V. Polypyrrole Nanoribbon Based Chemiresistive Immunosensors for Viral Plant Pathogen Detection. Anal. Methods 2013, 5, 3497, doi:10.1039/c3ay40371h.

27. Aydemir, N.; Malmström, J.; Travas-Sejdic, J. Conducting Polymer Based Electrochemical Biosensors. Phys. Chem. Chem. Phys. 2016, 18, 8264–8277, doi:10.1039/C5CP06830D.

28. Rashid, J.I.A.; Yusof, N.A. The Strategies of DNA Immobilization and Hybridization Detection Mechanism in the Construction of Electrochemical DNA Sensor: A Review. Sens. Bio-Sens. Res. 2017, 16, 19–31, doi:10.1016/j.sbsr.2017.09.001.

29. Shirale, D.J.; Bangar, M.A.; Park, M.; Yates, M.V.; Chen, W.; Myung, N.V.; Mulchandani, A. Label-Free Chemiresistive Immunosensors for Viruses. Environ. Sci. Technol. 2010, 44, 9030–9035, doi:10.1021/es102129d.

30. Dai, Y.; Liu, C.C. Recent Advances on Electrochemical Biosensing Strategies toward Universal Point-of-Care Systems. Angew. Chem. Int. Ed. 2019, 58, 12355–12368, doi:10.1002/anie.201901879.

31. Magar, H.S.; Hassan, R.Y.A.; Mulchandani, A. Electrochemical Impedance Spectroscopy (EIS): Principles, Construction, and Biosensing Applications. Sensors 2021, 21, 6578, doi:10.3390/s21196578.

32. Yang, C.; Jadhav, S.R.; Worden, R.M.; Mason, A.J. Compact Low-Power Impedance-to-Digital Converter for Sensor Array Microsystems. IEEE J. Solid-State Circuits 2009, 44, 2844–2855, doi:10.1109/JSSC.2009.2028054.

33. Stelzle, M.; Weissmueller, G.; Sackmann, E. On the Application of Supported Bilayers as Receptive Layers for Biosensors with Electrical Detection. J. Phys. Chem. 1993, 97, 2974–2981, doi:10.1021/j100114a025.

34. Labib, M.; Sargent, E.H.; Kelley, S.O. Electrochemical Methods for the Analysis of Clinically Relevant Biomolecules. Chem. Rev. 2016, 116, 9001–9090, doi:10.1021/acs.chemrev.6b00220.

35. Bontidean, I.; Berggren, C.; Johansson, G.; Csöregi, E.; Mattiasson, B.; Lloyd, J.R.; Jakeman, K.J.; Brown, N.L. Detection of Heavy Metal Ions at Femtomolar Levels Using Protein-Based Biosensors. Anal. Chem. 1998, 70, 4162–4169, doi:10.1021/ac9803636.

36. Heine, V.; Kremers, T.; Menzel, N.; Schnakenberg, U.; Elling, L. Electrochemical Impedance Spectroscopy Biosensor Enabling Kinetic Monitoring of Fucosyltransferase Activity. ACS Sens. 2021, 6, 1003–1011, doi:10.1021/acssensors.0c02206.

37. Felix, F.S.; Angnes, L. Electrochemical Immunosensors – A Powerful Tool for Analytical Applications. Biosens. Bioelectron. 2018, 102, 470–478, doi:10.1016/j.bios.2017.11.029.

38. Chen, C.; Jiang, Y.; Kan, J. A Noninterference Polypyrrole Glucose Biosensor. Biosens. Bioelectron. 2006, 22, 639–643, doi:10.1016/j.bios.2006.01.023.

39. Jain, R.; Jadon, N.; Pawaiya, A. Polypyrrole Based next Generation Electrochemical Sensors and Biosensors: A Review. TrAC Trends Anal. Chem. 2017, 97, 363–373, doi:10.1016/j.trac.2017.10.009.

40. Maas, M.N.; Hintzen, J.C.J.; Löffler, P.M.G.; Mecinović, J. Targeting SARS-CoV-2 Spike Protein by Stapled HACE2 Peptides. Chem. Commun. 2021, 57, 3283–3286, doi:10.1039/D0CC08387A.

41. McMaster, W.C.; Kouzelos, J.; Liddle, S.; Waugh, T.R. Tendon Grafting with Glutaraldehyde Fixed Material. J. Biomed. Mater. Res. 1976, 10, 259–271, doi:10.1002/jbm.820100207.

42. Habeeb, A.F.S.A.; Hiramoto, R. Reaction of Proteins with Glutaraldehyde. Arch. Biochem. Biophys. 1968, 126, 16–26, doi:10.1016/0003-9861(68)90554-7.

43. Letko, M.; Marzi, A.; Munster, V. Functional Assessment of Cell Entry and Receptor Usage for SARS-CoV-2 and Other Lineage B Betacoronaviruses. Nat. Microbiol. 2020, 5, 562–569, doi:10.1038/s41564-020-0688-y.

44. Shang, J.; Wan, Y.; Luo, C.; Ye, G.; Geng, Q.; Auerbach, A.; Li, F. Cell Entry Mechanisms of SARS-CoV-2. Proc. Natl. Acad. Sci. 2020, 117, 11727–11734, doi:10.1073/pnas.2003138117.

45. Zhang, H.; Penninger, J.M.; Li, Y.; Zhong, N.; Slutsky, A.S. Angiotensin-Converting Enzyme 2 (ACE2) as a SARS-CoV-2 Receptor: Molecular Mechanisms and Potential Therapeutic Target. Intensive Care Med. 2020, 46, 586–590, doi:10.1007/s00134-020-05985-9.

46. Lupala, C.S.; Li, X.; Lei, J.; Chen, H.; Qi, J.; Liu, H.; Su, X. Computational Simulations Reveal the Binding Dynamics between Human ACE2 and the Receptor Binding Domain of SARS-CoV-2 Spike Protein; Biophysics, 2020;

47. Lupala, C.S.; Kumar, V.; Li, X.; Su, X.; Liu, H. Computational Analysis on the ACE2-Derived Peptides for Neutralizing the ACE2 Binding to the Spike Protein of SARS-CoV-2; Biophysics, 2020;

48. Nevola, L.; Giralt, E. Modulating Protein–Protein Interactions: The Potential of Peptides. Chem. Commun. 2015, 51, 3302–3315, doi:10.1039/C4CC08565E.

49. Bigdeli, I.K.; Yeganeh, M.; Shoushtari, M.T.; Zadeh, M.K. Electrochemical Impedance Spectroscopy (EIS) for Biosensing. In Nanosensors for Smart Manufacturing; Elsevier, 2021; pp. 533–554 ISBN 978-0-12-823358-0.

50. Do, J.-S.; Wang, S.-H. On the Sensitivity of Conductimetric Acetone Gas Sensor Based on Polypyrrole and Polyaniline Conducting Polymers. Sens. Actuators B Chem. 2013, 185, 39–46, doi:10.1016/j.snb.2013.04.080.

51. Zhao, C.; Wang, H.; Zhang, H. Bio-Inspired Artificial Ion Channels: From Physical to Chemical Gating. Mater. Chem. Front. 2021, 5, 4059–4072, doi:10.1039/D1QM00070E.

52. Rohatgi, V.K.; Saleh, A.K.M.E.; Rohatgi, V.K. An Introduction to Probability and Statistics; Wiley series in probability and statistics; 2nd ed.; Wiley: New York, 2001; ISBN 978-0-471-34846-7.

53. Czibere, L.; Burggraf, S.; Becker, M.; Durner, J.; Draenert, M.E. Verification of Lateral Flow Antigen Tests for SARS-CoV-2 by QPCR Directly from the Test Device. Dent. Mater. 2022, 38, e155–e159, doi:10.1016/j.dental.2022.03.005.

54. Bekliz, M.; Adea, K.; Puhach, O.; Perez-Rodriguez, F.; Marques Melancia, S.; Baggio, S.; Corvaglia, A.-R.; Jacquerioz, F.; Alvarez, C.; Essaidi-Laziosi, M.; et al. Analytical Sensitivity of Eight Different SARS-CoV-2 Antigen-Detecting Rapid Tests for Omicron-BA.1 Variant. Microbiol. Spectr. 2022, 10, e00853–22, doi:10.1128/spectrum.00853-22.

55. Deerain, J.; Druce, J.; Tran, T.; Batty, M.; Yoga, Y.; Fennell, M.; Dwyer, D.E.; Kok, J.; Williamson, D.A. Assessment of the Analytical Sensitivity of 10 Lateral Flow Devices against the SARS-CoV-2 Omicron Variant. J. Clin. Microbiol. 2022, 60, e02479–21, doi:10.1128/jcm.02479-21.

56. Walsh, K.A.; Jordan, K.; Clyne, B.; Rohde, D.; Drummond, L.; Byrne, P.; Ahern, S.; Carty, P.G.; O’Brien, K.K.; O’Murchu, E.; et al. SARS-CoV-2 Detection, Viral Load and Infectivity over the Course of an Infection. J. Infect. 2020, 81, 357–371, doi:10.1016/j.jinf.2020.06.067.

57. Corman, V.M.; Haage, V.C.; Bleicker, T.; Schmidt, M.L.; Mühlemann, B.; Zuchowski, M.; Jo, W.K.; Tscheak, P.; Möncke-Buchner, E.; Müller, M.A.; et al. Comparison of Seven Commercial SARS-CoV-2 Rapid Point-of-Care Antigen Tests: A Single-Centre Laboratory Evaluation Study. Lancet Microbe 2021, 2, e311–e319, doi:10.1016/S2666-5247(21)00056-2.

58. He, R.; Liu, H.; Niu, Y.; Zhang, H.; Genin, G.M.; Xu, F. Flexible Miniaturized Sensor Technologies for Long-Term Physiological Monitoring. Npj Flex. Electron. 2022, 6, 20, doi:10.1038/s41528-022-00146-y.

59. Zhao, Y.; Wang, B.; Hojaiji, H.; Wang, Z.; Lin, S.; Yeung, C.; Lin, H.; Nguyen, P.; Chiu, K.; Salahi, K.; et al. A Wearable Freestanding Electrochemical Sensing System. Sci. Adv. 2020, 6, eaaz0007, doi:10.1126/sciadv.aaz0007.

60. Aliofkhazraei, M.; Ali, N. Recent Developments in Miniaturization of Sensor Technologies and Their Applications. In Comprehensive Materials Processing; Elsevier, 2014; pp. 245–306 ISBN 978-0-08-096533-8.

61. Dahlin, A.B. Size Matters: Problems and Advantages Associated with Highly Miniaturized Sensors. Sensors 2012, 12, 3018–3036, doi:10.3390/s120303018.

