## supplemental figures for "Development and Analytical Evaluation of a Point-of-Care Electrochemical Biosensor for Rapid and Accurate SARS-CoV-2 Detection"

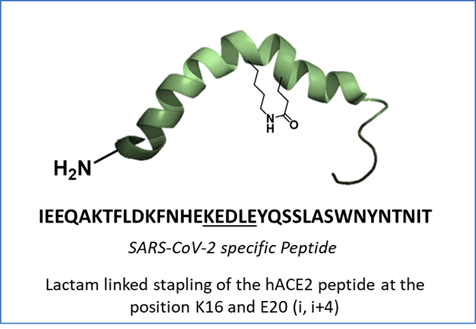

**Figure S1.** Lactam-stapled hACE-2 peptide sequence and design for functionalizing biosensor. The sequence and image were adopted from Maas et.al [40].

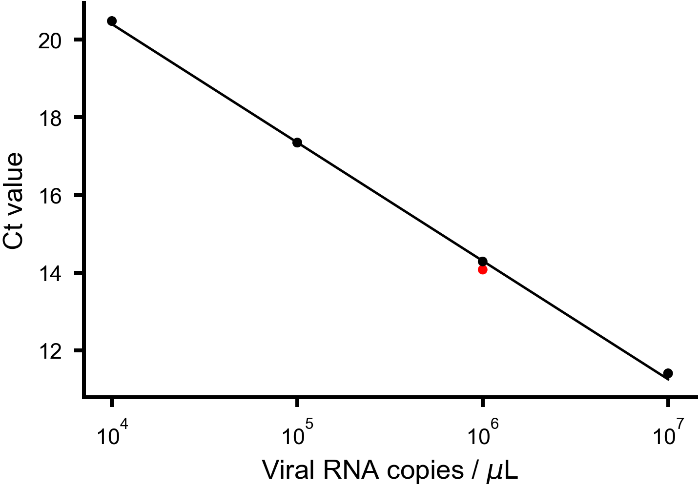
**Figure S2.** Standard curve generated for RT-PCR quantification of viral stocks. The standard curve is generated using the 10-fold serially diluted SARS-CoV-2 positive control of the known copy number provided with the Coronavirus COVID 19 Genesig real time PCR assay kit. The red dot on the curve indicates unknown sample concentration.

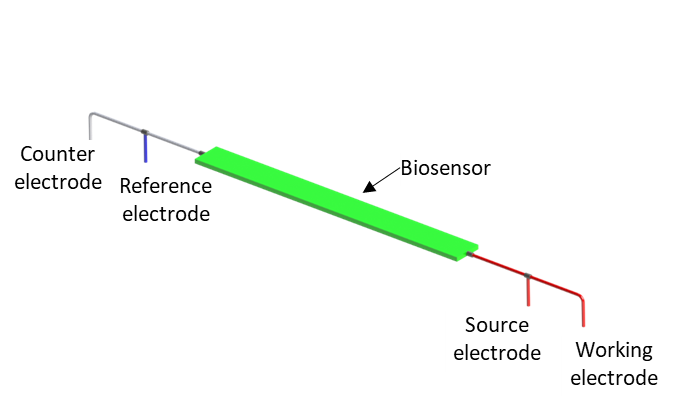

**Figure S3.** Connection configuration of potentiostat to the biosensor (green strip). To convert the standard set-up into a two-electrode system the working and source electrodes are coupled together, while counter and reference electrodes are shorted.

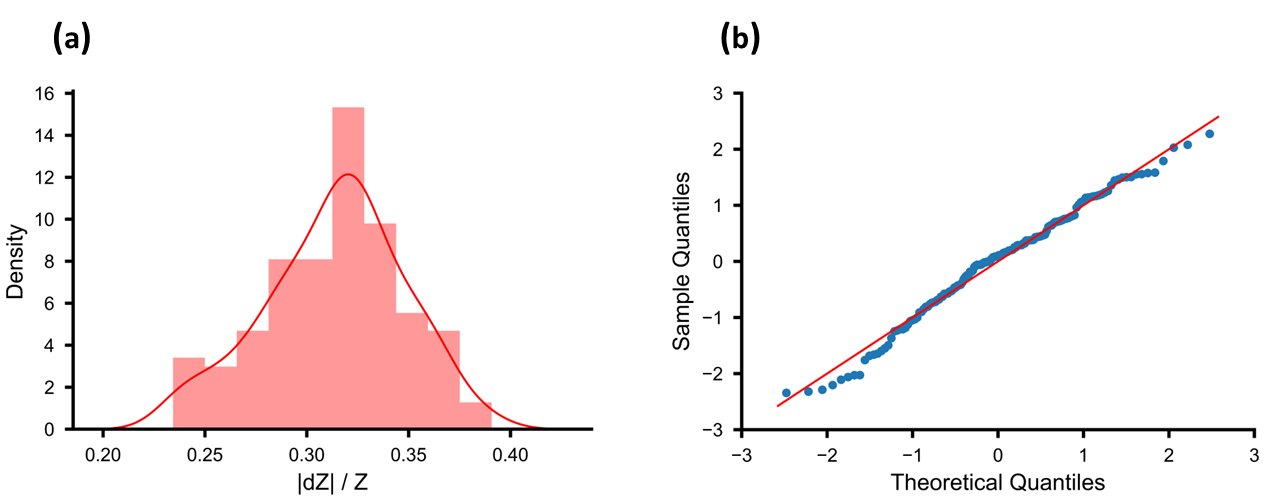

**Figure S4.** Normality test of distribution of relative impedance change of the biosensor. (a) Distribution of relative impedance (|dZ|/Z) change, (b) Q-Q plot of |dZ|/Z for 40 TCID50/mL from 50 independent experiments.

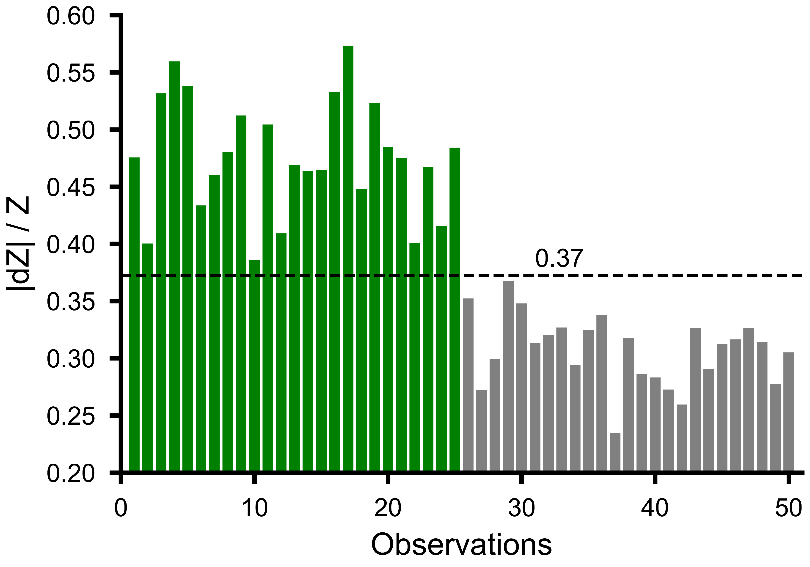

**Figure S5.** Threshold line generated using machine learning model for detection of virus. Y-axis indicates relative impedance value (|dZ|/Z) and X-axis represent independent tests. Green and grey bars represent media control and virus (4 TCID_50_/mL) respectively. The dotted black line represents threshold line for positive and negative classification of individual tests.

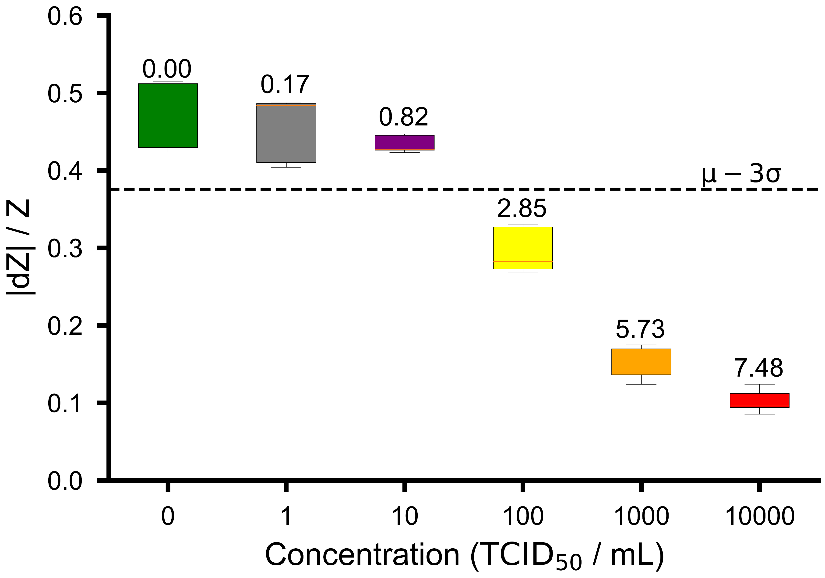

**Figure S6.** Assessment of Limit of Detection (LoD) through 10-fold Dilution Experiments. The figure illustrates the relative impedance change (|dZ|/Z) observed for various virus concentrations and a media control. The media control is highlighted in the green box, while different virus concentrations are depicted by boxes in shades of Grey, Purple, Yellow, Orange, and Red, representing viral RNA copies of 0.1 TCID_50_/mL, 1 TCID_50_/mL, 10 TCID_50_/mL, 100 TCID_50_/mL, and 1000 TCID_50_/mL, respectively. The corresponding sf values are provided as annotations above each box. A black dashed line signifies 3 standard deviations below the mean of relative impedance change from the control data. The red line within each box indicates the median value associated with that concentration.

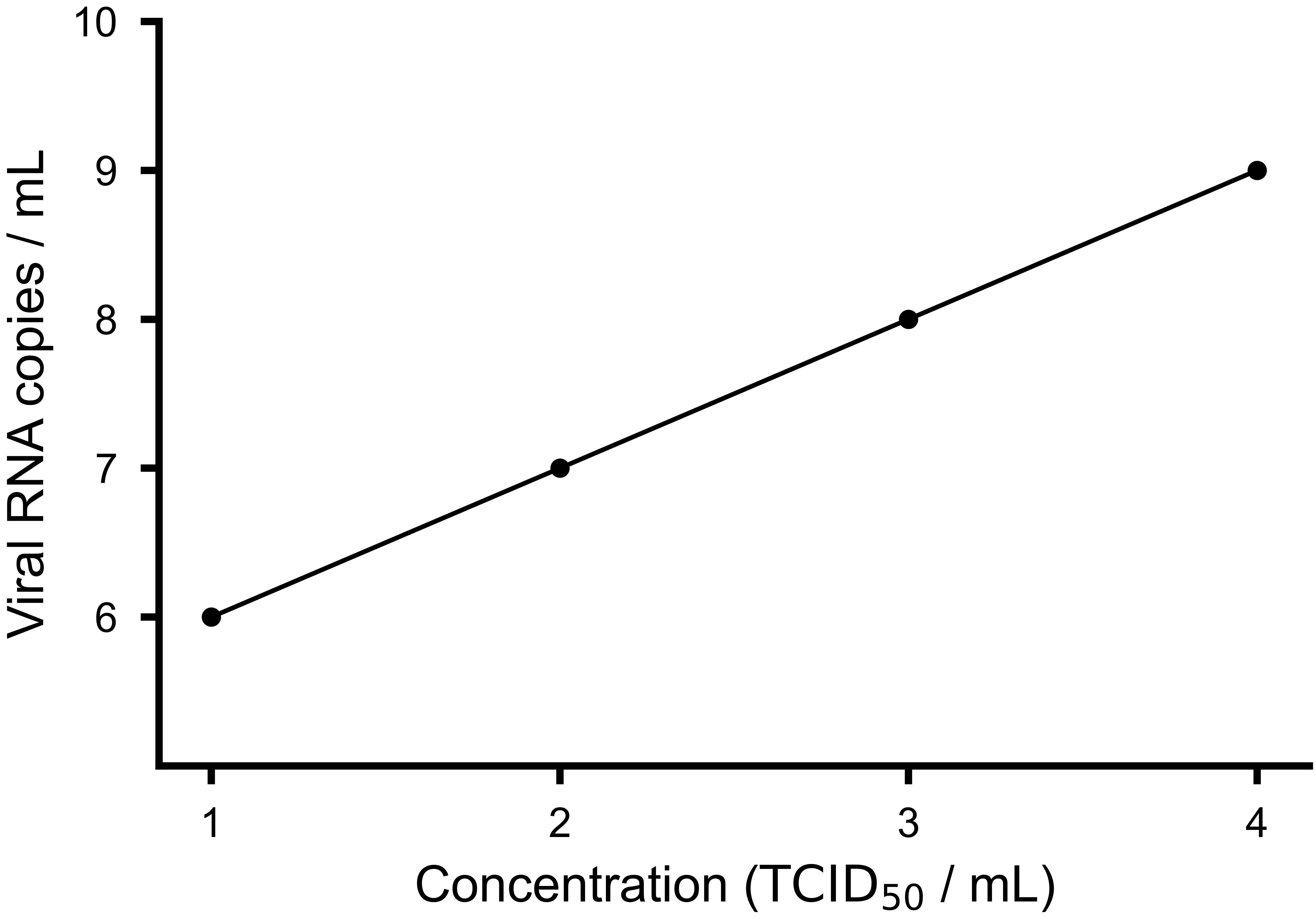

**Figure S7:** Correlation of TCID_50_/mL and viral RNA copies/mL for SARS-CoV-2 Delta and Omicron. Both the axes are in log_10_ scale.

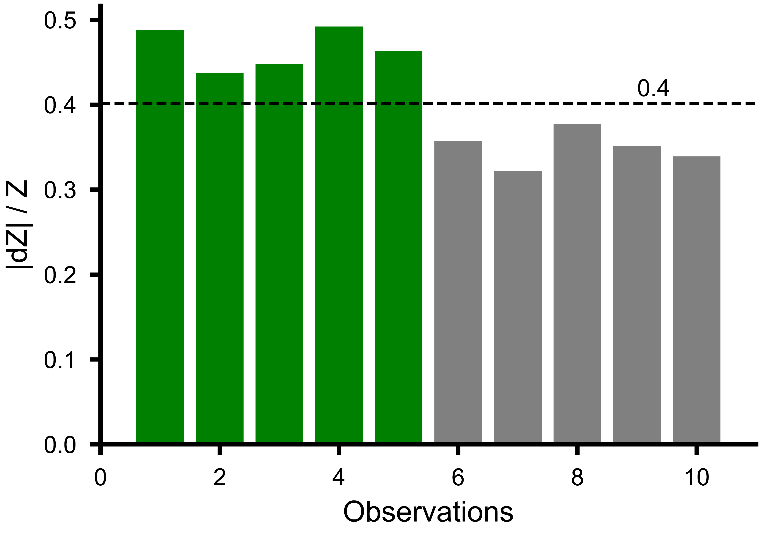

**Figure S8.** Response of delta variant virus- y-axis indicates relative impedance value (|dZ|/Z). Green, and grey bars represent media control and virus (4 TCID_50_/mL) respectively. The dotted black line represents threshold line for positive and negative classification of individual tests.
